# Consciousness in Non-REM-parasomnia episodes

**DOI:** 10.1101/2024.06.02.597000

**Authors:** Francesca Siclari

## Abstract

Sleepwalking and related parasomnias are thought to result from incomplete awakenings out of Non rapid eye movement (Non-REM) sleep. Non-REM parasomnia behaviors have been described as unconscious and automatic, or related to vivid, dream-like conscious experiences. Similarly, some observations have suggested that patients are unresponsive during episodes, while others that they can interact with their surroundings. To better grasp and characterize the full spectrum of consciousness and environmental disconnection associated with behavioral episodes, 35 adult patients with Non-REM sleep parasomnias were interviewed in-depth about their experiences. The level of consciousness during parasomnia episodes was reported to be variable both within and between individuals, ranging from minimal or absent consciousness and largely automatic behaviors (frequently/always present in 36% of patients) to preserved conscious experiences characterized by delusional thinking of varying degrees of specificity (65%), often about impending danger, variably formed, uni- or multisensory hallucinations (53%), impaired insight (77%), negative emotions (75%) and variable, but often pronounced amnesia (30%). Patients described their experiences as a dream scene during which they felt awake (‘awake dreaming’). Surroundings were either realistically perceived, misinterpreted (in the form of perceptual illusions or misidentifications of people) or entirely hallucinated as a function of the prevailing delusion. These observations suggest that the level of consciousness and sensory disconnection in Non-REM parasomnias is variable and graded. In their full-fledged expression, Non-REM parasomnia experiences feature several core features of dreams. They therefore represent a valuable model for the study of consciousness, sleep-related sensory disconnection and dreaming.

## Introduction

Sleepwalking (somnambulism) and related parasomnias refer to abnormal awakenings out of Non-rapid eye movement (Non-REM) sleep^1^, during which individuals may interact with their surroundings in a state of altered consciousness^2–4^. Parasomnia episodes can manifest as brief and simple actions, like sitting up and talking, or more complex behaviors, including engaging in conversations, leaving the bed, or manipulating objects. In extreme cases, sleepwalkers have been reported to drive, commit sexual assault, homicide and pseudo-suicide, resulting in personal tragedies and legal complexities^5–7^. Although rare, these incidents prompt fundamental inquiries into the state of consciousness of sleepwalkers and the nature of their experiences during parasomnia episodes. Already in 1878, Yellowlees convincingly described the dramatic case of a 28-year-old sleepwalker who attacked an imaginary white beast in his dream, only to wake up and realize that instead, he had just fatally injured his 18-month-old baby^5^. He suggested to name this condition ‘somnomania’, to distinguish it from other episodes of somnambulism. However, with the advent of the first electroencephalographic (EEG) recordings of somnambulistic episodes in the 1960ies, the idea that sleepwalking could be related to dreaming lost support. Somehow unexpectedly, sleepwalking was found to occur out of slow wave sleep, a stage that especially at the time was considered largely dreamless, in stark opposition to the recently discovered rapid eye movement (REM) sleep^1^. Although later, it was acknowledged that ‘apparent dreams’ could be recalled by sleepwalkers, these reports were judged unreliable, not necessarily related to the previous sleep period and possibly confabulated during the awakening^8^. In a famous case of somnambulistic homicide, it was even argued that Non-REM parasomnia behaviors are ‘preprogrammed’, precluding the possibility to form or execute conscious intents, similar to other forms of automatisms seen for instance in epileptic seizures^6^. Yet, a study in which patients with sleep terrors were interviewed immediately after parasomnia episodes, found that recall of mental content was surprisingly frequent^9^ (58%), consisting in terrifying scenes in which individuals were in danger of being crushed, enclosed by walls, or attacked by enemies. In the following years, paralleling the observation that dreaming could also occur in Non-REM sleep (reviewed in^10^), the idea that sleepwalking episodes could reflect dream-related behaviors, as opposed to largely unconscious ambulatory automatisms, gained support. Several case studies reported on patients with sleepwalking and sleep terrors who had vivid experiences, most often involving threatening, dream-like scenes^11–15^, but also visual hallucinations^16–18^, delusions^19–21^ and behavior reflecting the ‘dream’ content^17,18,22^. Studies in which the recall of conscious experiences was retrospectively quantified and described in detail revealed that 66 to 91% of adult patients with sleepwalking and/or sleep terror remembered at least one episode associated with mental activity over their lifetime ^13,15,18^, while this proportion was much lower in children (around 33%)^21^. These contrasting reports, of dream-related behaviors and interactions with the environment on the one hand, and unconscious automatisms with environmental unresponsiveness on the other hand, raise several questions: can behavior during parasomnia episodes truly occur in the absence of consciousness, as initially assumed, or do affected individuals always experience something, and then simply forget? Is consciousness associated with parasomnia episodes an all-or-none phenomenon, or is it graded? Do sleepwalkers perceive their environment, and how exactly?

In the present paper, answers to these questions were sought by interviewing 35 patients with Non-REM parasomnia in depth. Instead on focusing exclusively on whether patients ‘dreamt’ during the episodes or not, this work aimed to outline, in as much detail as possible, the whole spectrum of consciousness associated with parasomnia episodes, both with regards to the level of consciousness as well as its contents. Qualitative features of conscious experiences associated with NREM parasomnia episodes were classified in broad cognitive categories (consciousness, memory, thinking, perception, metacognition and emotions).

## Methods

### Patients

35 adult patients with a diagnosis of disorder of arousal (confusional arousals, sleepwalking and/or sleep terrors) underwent a semi-structured interview by the author about their experiences during parasomnia episodes. Patients were recruited at the outpatient Center for Investigation and Research on Sleep at the Lausanne University Hospital, Switzerland, or by word of mouth, as part of a larger study comprising high-density EEG sleep recordings^36^. The inclusion period ranged between June 2016 and August 2022. The diagnosis of disorder of arousal was made by the author of the study, a board-certified neurologist and sleep specialist, according to international criteria^11^ and after a clinical evaluation comprising an overnight polysomnography (PSG). Written informed consent was obtained by all the participants and the study was approved by the local ethics committee (commission cantonale éthique de la recherche sur l’être humain du canton de Vaud). All patients who agreed to participate in the study were interviewed, regardless of whether they remembered their parasomnia episodes or not. Patients in whom parasomnia episodes could not be clearly distinguished from sleep-related seizures, or those presenting evidence, during polysomnography, for concomitant REM sleep behavior disorder or REM sleep without atonia were not included in the present study. Patients presenting exclusively with sleep-related eating disorder or sexsomnia were also not included.

### Procedure

The author of the study conducted a semi-structured interview, focusing on the following aspects of parasomnia episodes: 1) *Memory* (recollection of experiences), 2) *Thinking* (ideas, beliefs held during episodes), 3) *Perceptual aspects* (presence of hallucinations or illusions, perception of the real environment), 4) *Metacognition* (insight, control over the experience), 5) *Behavior* (purposeful vs. automatic behavior), 6) *Emotions* (negative, positive, neutral) and 8) *other aspects* (including length and setting of the experience). For each of these domains, patients had to rate the occurrence on a 5-item scale: 1=never (applies to none of the episodes/experiences), 2=rarely (applies to less than half of episodes/experiences), 3=sometimes (applies to approximately half of the episodes/experiences), 4=often (applies to more than half of episodes/experiences) and 5=always (applies to all the episodes/experiences). In addition, patients were asked to detail their answers and to provide examples for each aspect. Some aspects that were not systematically addressed by the questions but consistently emerged from the interviews were also included as results and quantified whenever possible. Interviews were audiotaped and later transcribed. Examples of reports included in this publication were translated from French to English by the author.

## Results

### Patient characteristics

35 patients were interviewed [19 females, age 28.9 ± 8.2 yrs (average ± SD), range 18.3–56.4]. Age of onset of parasomnia episodes was 8.6 ± 4.5 yrs (range 3-18). Self-reported frequency of parasomnia episodes at the time of interview was distributed as follows: less than once a month: 5 patients (14%); ⁓once a month: 4 patients (12%), 2-3 times a month: 10 patients (29 %); ⁓once a week: 6 patients (17%); 2-3 times a week 5 patients (14%) and almost every night: 5 patients (14%). All patients had a history of confusional arousals, 24 patients (68%), had, in addition, a history of *both* sleep terrors and sleepwalking, 9 (25%) patients only of sleepwalking and one patient (3%) only of sleep terrors. Since almost all patients presented several types of parasomnia episodes and the distinction between the different types is mainly qualitative and not straightforward to make in all cases, all patients and episodes were considered together, regardless of the type of parasomnia. Eight patients had neurological or psychiatric comorbidities including migraine (n=2), multiple sclerosis (n=2), idiopathic hypersomnia (n=1), a history of mental anorexia (n=1) and a history of depression (n=2). None of the patients had ever experienced delusions or hallucinations unrelated to their parasomnia episodes.

### Characteristics of parasomnia episodes

The majority of patients (94%) could report what they had experienced during at least one parasomnia episode, and roughly one third (32%) estimated that they always or often remembered their experiences (Fig. 1A). The characteristics of these experiences are outlined in the next section. A description of parasomnia episodes with no or minimal experience (automatic behavior) and amnesia is provided at the end of the results section. So as not to distract from results, actual reports of experiences are provided in a separate table (Table 1).

**Figure 1.**
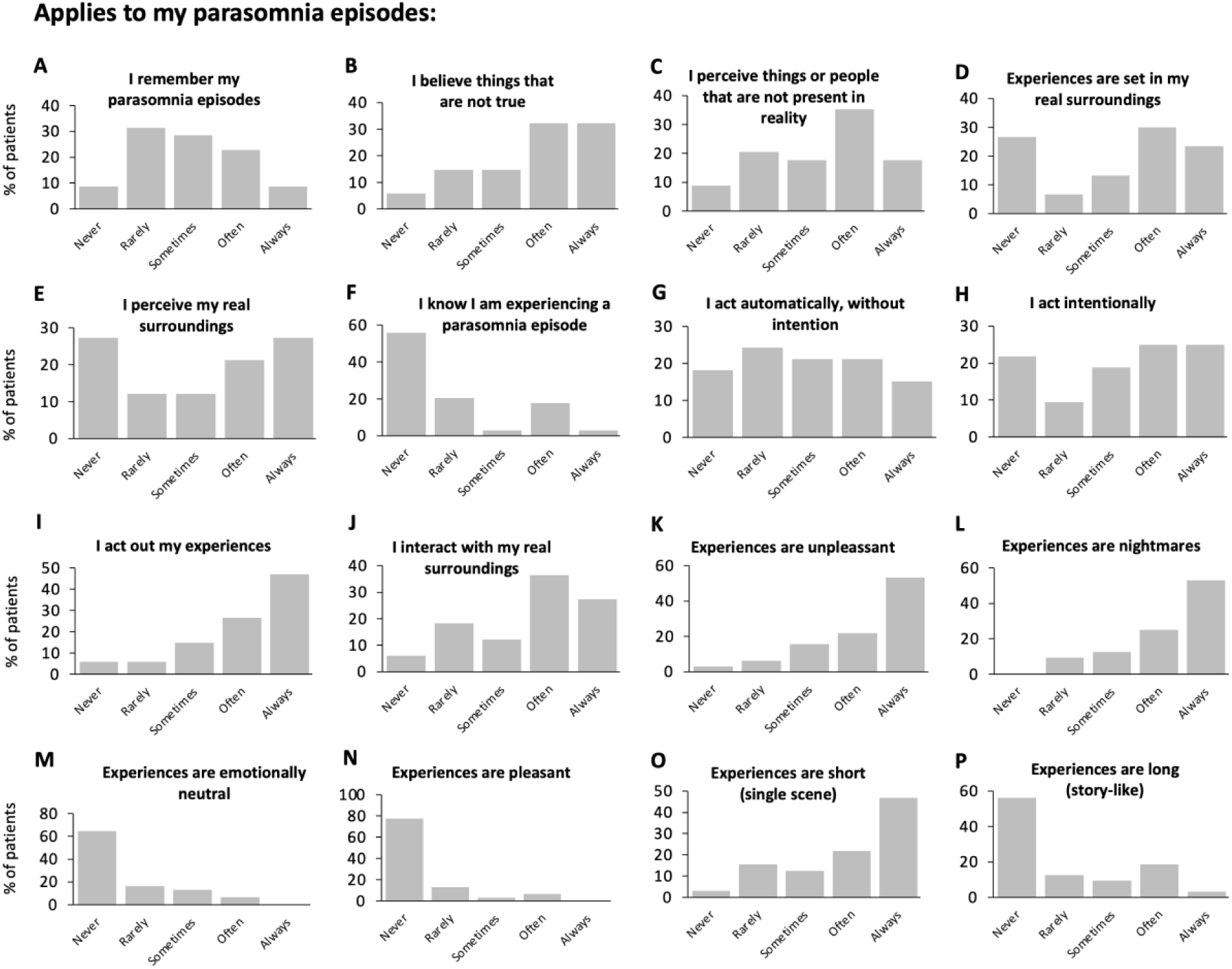
Self-reported frequency of parasomnia episode characteristics (n=35 patients). Patients rated the occurrence of each feature on 5-point scale: 1=never (applies to none of the episodes/experiences), 2=rarely (applies to less than half of episodes/experiences), 3=sometimes (applies to approximately half of the episodes/experiences), 4=often (applies to more than half of episodes/experiences) and 5=always (applies to all the episodes/experiences).

**Table 1:** Examples of reports of parasomnia experiences for different cognitive domains.

| Number | Report | Comment |
| --- | --- | --- |
| **Delusional thinking** |  |  |
| 1 | I wake up suddenly and I know that I am in danger of dying. It is not more concrete than that. There are no images. | Simple delusional thinking. |
| 2 | Once I woke up and knew a bomb was going to go off in my house. I woke up all my children and took them outside. We waited outside for the bomb to go off. | More specific delusional thinking. |
| 3 | I wake up in my bed next to my husband. A strong voice inside me, associated with a black mass, tells me I have to go somewhere to serve my sentence, to be punished for something I did (I don't know what). The punishment would have been less painful if I had taken my old nightmares more seriously. I ask my husband to help me, but he doesn't wake up. I panic because if I comply with this voice's instructions, I am going to sacrifice myself and lose everything, and will not see my children again. If I don't obey, things will get worse (I don't know how exactly). | Complex and specific delusional thinking. |
| 4 | I wake up and wonder if my daughter has put on her hydrocortisone cream, and then I set out to verify. It doesn't occur to me that she has stopped this treatment a long time ago. | Delusion that is specific to the patient's (former) context. |
| 5 | You see, it is like when you forward a movie with a remote control, image by image. The last image, just before death, is when I scream, that's my last chance to save my skin. In the next frame, I would have been squashed, riddled with bullets, or I would have fallen off the cliff. It is as if I anticipated the next image, the next step. This anticipation of what is to come is very clear, it is more real than in real life. | Report illustrating the impending nature of the imaginary threat. |
| **Hallucinations** |  |  |
| 6 | I am terrified. I know there is someone in the room who wants to harm me. Sometimes I see this person clearly next to my bed, sometimes I just sense the person behind me. I know that if I don't do something I will die, so I start to run, I look for an exit. | Variably formed hallucinations (ranging from sensed presence to visually vivid hallucinations). |
| 7 | I often dream about people who want to hurt me. Sometimes they are behind me. Sometimes I see them. I can usually tell whether it is a woman or a man. When I take clonazepam, their characteristics are much more vague. | Variably formed hallucinations. |
| 8 | A few years ago, I regularly saw a bearded man on my right. Currently I see a shadow that passes from one side to the other. | Formed and unformed visual hallucinations. |
| 9 | I once dreamt of a little dog that was dying of thirst because it was too hot. I can still picture this dog lying on its side, next to my radiator, his fur humid from the perspiration. | Visually vivid, formed and detailed hallucination. |
| 10 | I remember seeing a spider, the image I have is the one of a big blue tropical spider, very clear, perfectly illuminated. | Visually vivid, formed hallucination. |
| **Perception of the real environment, illusions, misidentifications** |  |  |
| 11 | I can see images relatively clearly, as if the light was on. However, when I wake up, I realize that everything is dark. | Example showing how some aspects of the real environment are hallucinated or misinterpreted. |
| 12 | During the first seconds of my awakening, the whole luminous intensity created by my mind vanishes. | Example showing how some aspects of the real environment are hallucinated or misinterpreted. |
| 13 | I tend to get hurt during my episodes when I think that I am somewhere else than in my real surroundings. Sometimes I dream that I am at work, and I hurt my head against the pitched roof of my bedroom because in my head, I am at work. As if I saw that as a mental image, like a big memory of what I know from being awake. | Example of how the hallucinated environment may lead to accidents. |
| 14 | Once in the news I saw a video of a kid at a kite convention who got carried away by a huge kite, he rose several meters above the ground but managed to hold on to the kite and fortunately was not harmed. Sometime later I dreamt of this scene; in my dream I was in the park and saw this kid. Suddenly the kid [makes a whooshing sound] takes off, I run after him to try and catch him, but then I hit the wall. The real wall. | Example of how the hallucinated environment may lead to accidents. Also shows how behavior can be triggered by a surprising event in a dream. |
| 15 | Once I saw a window in front of my bed, and through this window I could see a tree and a person observing me. Then I woke up and saw a gradual transition between the image of the window and the wall of my room. Like a 'fade' effect on a power point presentation. | Transition between perceiving the hallucinatory and the real environment. |
| 16 | I dreamt that I was driving and that the breaks of the car did not work, so I opened the door of the car and tried to slow down by touching the ground with my left foot. My mother found me sitting on the edge of the bed, screaming, with my left leg violently hitting the floor. When she switched on the lights, the interior of the car and the street in front of me immediately dissolved, as if someone had 'peeled off' a thin layer from my surroundings, to unmask a different reality. | Transition between perceiving the hallucinatory and the real environment. |
| 17 | Once during a sleep terror, I dreamt that someone caught me with a fishnet. When I woke up, I realized that I had hit a coat hanger while running, and that what I had thought was a fishnet were in fact a couple of jackets that had fallen on me. | Perceptual illusion. |
| 18 | Next to my bed there was a drying rack with clothes on it. During an episode I made up a theory that they were power lines with bats on them, and I tried to pass under the rack while protecting myself with my blanket. Then my mother came in and asked me what I was doing. I told her that these were power lines, and that she had to be very careful not to touch them. | Perceptual illusion. |
| 19 | Things are really well done in those dreams. The table in my room for instance, will not be a table but will have the same shape and dimensions as the table. But then sometimes this is not the case. And that's when I get hurt. Once for instance I dreamt that I was in a prairie, and that an old wheel of a windmill was rolling towards me. In my dream I started to lean against a barrier, but in reality there was nothing, so I lost my balance and fell off my bunk bed, on the tiled floor. I suffered a concussion and a contusion of my shoulder. | Account of perceptual illusions and hallucinations, and how the latter can lead to accidents. |
| 20 | At the beginning of my relationships with men, I sometimes wake up and have the impression that someone evil is lying next to me. My boyfriends try to be nice and to reassure me, they talk to me and touch my arm. But I don't see their real face, only the face of the evil man, sometimes even only a silhouette or a dark shadow. It is very difficult to realize that this not real, because the person next to me is really talking to me and touching me. That's when I become dangerous because I try to defend myself. | Example of how misidentifications can lead to violent behaviors. And how hallucinations can occur in one sensory modality (visual) and perception of the real environment in another (tactile, auditory). |
| 21 | Once I dreamt that I was talking to my mother, I was showing her a box with lots of things in it. Then I woke up and realized that everything was dark and that instead of holding a box, I was holding my pillow. | Co-existence of hallucinations and perceptual illusions. |
| 22 | If there is no light, there is this virtual veil in front of my eyes, I am in my imaginary world, I walk into walls and stumble upon things. But if there is light, then it is different. Once at the army I slept with the lights on, I got up and started to go through the things in my bag. My comrade asked me what I was doing, and I answered that there were snakes. He said 'No, these are the straps of your bag', and apparently I protested. | Example of how ambient illumination may favor illusions over hallucinations. |
| **Metacognition: impaired insight, ambivalence** |  |  |
| 23 | I often fight with my girlfriend during the episodes. She tells me I am having one of my episodes and gives me instructions on what to do. I tell myself she has a point because it is true that I have had sleepwalking episodes in the past. Still, I don't agree, I get angry, because for me this is not an episode, it's reality. It is hard for me to hear that it is an episode when I really see things. That's why I can get really angry. | Example illustrating how false beliefs are difficult to challenge. |
| 24 | I recently told my boyfriend that he had spiders on his head. He told me I was dreaming, and that got me mad. I started protesting, and while I was speaking, I progressively realized that I was wrong. | Non-criticized hallucinations, with gradual recovery of insight. |
| 25 | Once during a sleep terror, I decided to jump out of the window, knowing that I would get hurt, because I estimated that the risk of dying by jumping out of the window was still lower than the risk of getting killed by the people who were after me. | Example of how fixity of beliefs can lead to risky behaviors. Patients can be perfectly aware of the risks they take. |
| 26 | I know that I am delirious, but I am too absorbed in the dream. | Ambivalence |
| 27 | I see things and believe in them, but at the same time I know that there is another world that is real. | Ambivalence |
| 28 | Sometimes during my episodes, it is as if I was talking to myself. I ask myself: 'T. [name of the patient], what is going on?' I answer: 'I am at work, do not disturb me'. I say to myself: 'No, you are in your bed', and then I reply: 'That's not true, stop it. I can see my colleagues, so it is not possible'. I can switch very rapidly between the two scenarios: at one moment I can touch my bed and feel that I am lying down, and the next moment I see my colleagues from work, and I also see myself wearing a uniform. | Ambivalence resulting in 'dissociative' inner dialogue. |
| 29 | Little by little I realize, but I have to say things several times. Slowly but surely, while repeating the same sentences over and over again, I find that they do not make sense anymore. | Gradual recovery of insight. |
| 30 | I suddenly wake up, and for the first seconds I still have this sensation of panic, I don't hallucinate but I still have the impression of being in danger, or that there are insects in my bed, even if I don't see them. Even though I know it was a dream, I still feel the need to get up, switch on the light and look at my bed to make sure there are no insects. Afterwards I feel silly for having verified. | Gradual recovery of insight. |
| 31 | I once dreamt that a colleague of mine slept at my house and that I gave him my jacket, because he was cold. Then I dreamt that he was leaving with the jacket on and my keys in the pocket, so I ran after him to recover the keys. When I arrived at the door, I saw my keys on a cabinet, and that's when I told myself that it must have been a dream. | Recovery of insight upon noticing discrepancies between real and imagined surroundings/scenario. |
| 32 | Once I dreamt that a burglar was hiding under the sofa. I left my bed, crawling on all fours, until I reached the living room. There, I hid behind a door to look under the sofa, but there was no one there. That woke me up. | Recovery of insight upon noticing discrepancies between. real and imagined surroundings/scenario. |
| **Purposeful behavior** |  |  |
| 33 | Once I went out on the balcony because an intruder was going to enter my bedroom. I held a chair above my head, ready to hit the intruder with it as soon as he would step out on the balcony. But nothing happened. I woke up because my arms started to tingle from holding them above my head. I felt silly standing naked on the balcony at 5 o' clock in the morning. | Description of purposeful behavior in relation to a delusion. |
| 34 | I dreamt that a dog was lying next to my bed. He was dying of thirst because of the heat, so I went to the kitchen, filled a bowl with water and placed it next to my bed. I also placed a kitchen towel under the bowl to prevent the water from splashing on the floor. | Description of purposeful behavior in relation to a delusion. Same episode/patient as report 9. |
| **Negative emotions** |  |  |
| 35 | It is like a horror movie, you cannot imagine. It cannot get any more real than that. There is this emotional anticipation of death. There is more to it than death, it is knowing that you will die, but that on top of that, you will die in a horrible way. It is submerging, like an interior apocalypse. | Intensity of negative emotions related to sleep terrors. |
| 36 | I have never experienced such intense fear during wakefulness, even when I was once attacked in the street. After this attack, I was offered psychological help, which I refused. I wanted to tell them that I was used to being scared, and that what I experience in my awake nightmares is a thousand times worse than this attack in the street. | Comparison between fear during sleep terrors and in wakefulness. |
| 37 | In real life I am not a chicken. Once I was about to be mugged and I was courageous enough to scream and scare the attackers away. Another time I prevented a man from setting a building on fire, I ran after him, confronted him, and later went to trial to testify. Both times I wasn't that scared. I wonder why it is that at night I am so scared because when I am awake I don't "lose it" like that. | Comparison between fear during sleep terrors and in wakefulness. |
| **Other: vigilance** |  |  |
| 38 | I wake up with my heart beating really fast, I am disoriented, and I never know if what I just experienced was real. Despite this stress, I fall asleep quickly. Sometimes I scream and the next second I fall asleep. | Dissociation between emotional intensity from vigilance. |
| **Amnesia** |  |  |
| 39 | I woke up as usual, without any particular memories of the night. When I went to the bathroom, I noticed that the very expensive anti-aging cream I had bought the day before had been opened and was almost empty. Only then I remembered that during the night I had gotten up to put on almost the totality of the cream on my face. That was absurd of course, considering the price of the cream. | Example of how initially forgotten memories can be retrieved with the help of cues. |
| **Episodes with unconscious, minimally or partially conscious behavior** |  |  |
| 40 | Once on a trip to Africa, I slept in a dorm with twenty other girls. One of the girls came to bed later and saw me sitting on the edge of the bed, eyes wide open, gesturing with my hands. She thought I was haunted and woke up the other girls. Everyone started to scream. It is only then that I realized what I was actually doing. But I have no idea why. | Example of automatic behavior. |
| 41 | Sometimes I can say words that have nothing to do with one another. Sometimes my sentences are not grammatically correct and there is no intention behind them. | Example of automatic behavior. |
| 42 | It feels hot in the bed since my partner joined me a short while ago. I decide to go to the bathroom – a pressing need. On my way there I see my stepdaughter, standing bare breasted in the door frame. The light is on in the corridor. She urges me to stop. I ask her on an aggressive and peremptory tone if she has lost her mind, since I do not know what she is talking about. I go back to bed. In bed, my partner tells me that I screamed again. I had not noticed that I had been screaming and I am unable to say what could have | Example showing how patients may experience some aspects of the episode but not others (written report provided by patient). Also illustrates the frequently reported |
|  | scared me. I just remember going to the bathroom and seeing my stepdaughter who had gotten up in a hurry when she heard my screams, to see if her mother was well. | impression of not having been asleep yet before the parasomnia episode. |

#### Parasomnia episodes associated with conscious experiences

##### 1) Thinking

Almost all patients (94%) reported having had false ideas or convictions during episodes at least once, and for roughly two third of patients (65%) such mentation often or always occurred (Fig. 1B). The convictions frequently had danger or threat as a theme. The belief could be more or less elaborate, ranging from simply thinking that ‘something was off’ or that ‘something bad’ was about to happen, to more specific convictions, for example that an intruder was located in a particular room of their house or behind the door, that they were being followed by enemies or that their close ones were in danger. Examples of relatively isolated erroneous beliefs held during episodes, with increasing specificity and complexity, are reported in Table 1 (reports 1-3) and Fig. S1.

The conviction could be isolated or occur in relation to a dream-like scenario with hallucinations, as detailed in the next section. Interestingly, some delusions were consistently reported by different patients, including persecutory thoughts (that someone was following them and/or wanted to harm them or their close ones), ideas of a phantom intruder (unwanted presence of a stranger in their home), parasitosis (that insects or other organisms had invaded their bed, more rarely that they were on their body), delusions of infidelity (that their spouse or partner was being unfaithful), beliefs around respiratory difficulties (having swallowed something and choking on it, being strangled or suffocated, drowning), being threatened by moving elements of their environment (walls closing them in, objects falling from the ceiling, landslides, vehicle moving towards them, inability to control a car while driving) or by other impending disasters (bomb about to explode). Interestingly, most of these dramatic scenarios are rather unlikely to occur in real life. Other, more likely and less life-threatening beliefs were also reported and could be more specific to the patient’s context and life situation, including for instance the impression that the house was too messy and needed to be tidied, or beliefs of spousal infidelity. Sometimes, instead of a conviction, patients reported doubts, for instance about whether specific household duties had been carried out, or whether a regular medication had been taken (Table 1, report 4). Typically, the threat or danger was impending, that is, patients found themselves in a situation in which they could still flee or defend themselves, or else do what was necessary to prevent harm, even if they had little time: a bomb for instance was just about to go off, or a ceiling was close to the point of collapsing. One patient described poignantly how during his sleep terrors, he had the impression of experiencing the ‘last scene’ before death (Table 1 report 4). Often, a specific delusion recurred in a given patient for a certain period of time (although often with some variation), and then changed.

##### 2) Perceptual aspects

Almost all patients (91%) reported at least one episode with false perceptions (hallucinations), and roughly half of patients (53%) always or often experienced them during episodes (Fig. 1C). Patients spontaneously described mainly hallucinations of the visual type, which could either be set in the patient’s current location (often or always in 53 %, generally the bedroom, Fig. 1D), or in an entirely different, either familiar or unknown environment, such as the sea, the bottom of a volcano, the forest, a laboratory, or their workplace. For 27% of patients, sensory experiences were never related to the current environment (Fig 1D).

Visual hallucinations were mostly formed, but similar to the false beliefs, which ranged from vague to highly specific, the degree to which hallucinations were formed could vary, even within subjects, with shadows and silhouettes at one end of the spectrum, and detailed, highly realistic imagery of objects, animals or people on the other end (Table 2 and Fig. S1). The feeling of a presence^23^ (sensed presence) was described as always or frequently occurring by 14% of patients and was reported to have occurred at least once by 30% of patients. Examples of detailed imagery given by patients included moving tentacles, electrical wires, splashing waves, clowns, alligators, crabs, a camera, a steam roller, the headlights of a car, a robotic leg made of steel, a combine harvester, and a horse trailer. Reports of visual hallucinations with varying degrees of detailedness are listed in table 1 (Table 1, reports 5-10).

**Table 2:**
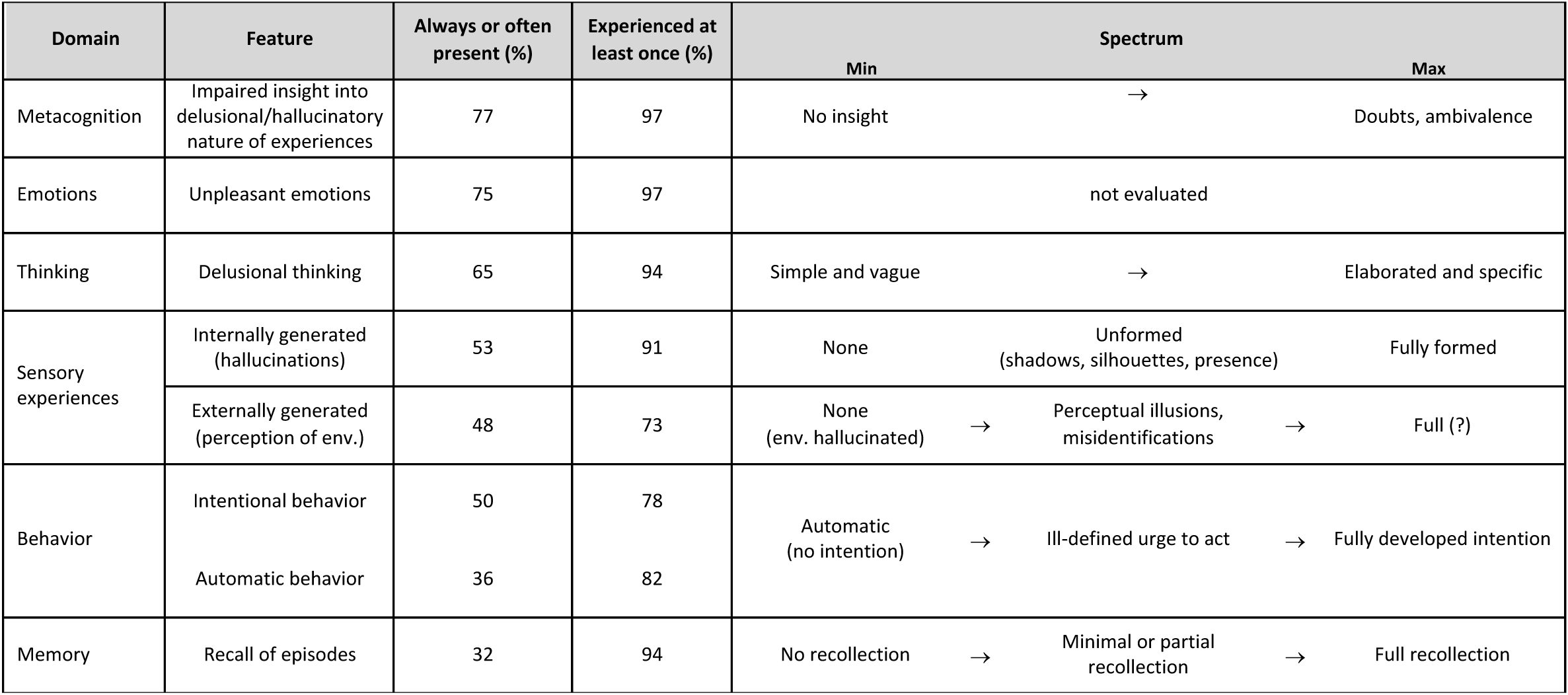
Summary of cognitive features of parasomnia experiences and their expression along a minimal to maximal gradient (spectrum).

When specifically asked about other sensory modalities, patients occasionally gave examples of auditory, tactile or vestibular hallucinations, like for instance a voice whispering or calling their name, an imaginary conversation with a person, the sound of a fictive alarm clock, physical touch when wrestling with enemies or intruders, or experiences of moving or falling.

##### 3) Perception of the real surroundings

Most patients (73%) reported having perceived elements of their real surroundings during parasomnia episodes at least once, and for roughly half of them (48%) this was often or always the case (Fig. 1E).

However, because oneiric scenarios are often set in the real surrounding of the patients, and patients are usually highly familiar with their bedrooms, it is sometimes difficult to know, from their accounts alone, to which degree they truly perceived their environment or whether they imagined it or parts of it, perhaps based on a good recollection. For instance, although it is generally dark in the bedroom, patients often report seeing objects in their surroundings “as if they were illuminated” and describe transitioning towards darkness only when they wake up, suggesting that some aspects of the purportedly perceived environment were actually hallucinated or misinterpreted (Table 1, reports 11 and 12). In other instances, the whole environment is entirely hallucinated. In these cases, patients move within fully imaginary surroundings, which bear no similarities to their real surroundings. Examples of such dream settings reported by patients include a river in the countryside, a forest at night, the patient’s classroom, a science laboratory, a landscape made of volcanic lava, the interior of a car that is about to plunge into a ditch, orchards and a campsite. Patients can hurt themselves or bedpartners during this type of episode because they collide with real elements, which are not part of their imaginary scenario, or because they lean on non-existing objects and lose their balance (Table 1, reports 13 and 14). Some patients could pinpoint the precise moment they transitioned to full wakefulness, when dream imagery was replaced by perceptions of their real surroundings, akin to a ‘fade’ effect in a power point presentation (Table 1, reports 15 and 16).

Apart from episodes in which the environment was fully hallucinated and others in which it appeared to be perceived as it is, there were instances in which patients *misinterpreted* environmental elements, that is, they experienced perceptual illusions, taking a real element for another (Table 1, reports 17 and 18). These perceptual illusions were most often in line with the prevailing delusional belief: one patient for instance removed the picture frames that were hanging on the wall next to her bed, believing she was picking apples, an activity that she had been doing during the previous day, and neatly arranged them next to her bed. Another patient dreamt that she was carrying dishes to her kitchen but woke up and realized she was carrying a cactus instead. Not infrequently, the imaginary objects display at least a crude resemblance to the real objects (Table 1, report 19).

Illusions may also involve people and in particular bedpartners who are believed to be someone else (misidentifications). For instance, during sleep terrors, patients may take well-meaning bedpartners or close ones for enemies who want to harm them. They may also recognize their family members as who they really are, but attribute malicious intentions to them. One young patient for instance recognized his father, but had the impression that the latter had been replaced by someone evil, as seen in Capgras syndrome^24^. From patients’ reports it becomes clear how in these cases, interactions with others can fuel the delusion and contribute to dangerous behaviors, as they reinforce the patients’ impression that what they experience is real (Table 1 report 20). Illusions may co-occur with hallucinations and with perceptions of the real environment (Table 1 report 21). Some reports suggest that the degree to which the environment is visually perceived vs. hallucinated or misinterpreted may depend on ambient illumination, with hallucinations occurring predominantly in the dark, and illusions or perception of the real environment when lights are on (Table 1 report 22).

##### 4) Metacognition

Patients do not usually criticize the beliefs or the hallucinations they experience during their episodes. Roughly half of them (56%) never realized that they were experiencing a parasomnia during the episode (Fig 1F). While some scenarios are not so implausible after all, most of them are, yet they are usually taken for real without a doubt. In fact, patients are so certain that what they experience corresponds to the reality that when they are confronted by their bedpartners about the possibility that they are dreaming or hallucinating, they often become angry and protest or argue (Table 1 examples 23 and 24). The unshakable certainty about the delusional thought also leads patients to act according to their convictions, sometimes at great risk. Patients may for instance leave the house or jump off windows to escape from impending disasters or imaginary pursuers, which can lead to accidents and injuries. Some patients are perfectly aware of the risks they take and of the potential consequences of their behavior (Table 1 report 25).

At times, usually towards the end of the episode, patients may experience doubts and ambivalence about whether what they believe or perceive is real. They may even consider the possibility that they are sleepwalking and ponder it against the impression that it all feels very real, “too real to be a dream” (Table 1 reports 26 and 27). This ambivalence, when it occurs, is invariably described as deeply disturbing. Patients feel caught between two possible interpretations of reality and try to find clues that help them decide on which one to adopt. One patient even reported experiencing a dissociative dialogue between these two opposing viewpoints (Table 1, report 28).

Lucidity with respect to the situation usually recovers gradually towards the end, when the belief the patient held during the episode ceases to make sense (Table 1 reports 29 and 30). Some patients eventually become lucid upon noticing discrepancies between the perceived and imaginary scenario (Table 1 reports 31 and 32).

##### 5) Behavior

The behavior that is displayed during the parasomnia episode can be either intentional and congruent with the false convictions and hallucinations or occur without apparent intentions and even awareness (see section ‘parasomnia episodes with no experiences or amnesia’ below). 40% of patients reported that they often or always acted intentionally and only rarely or never automatically, roughly a fourth (27%) that they sometimes acted intentionally and sometimes automatically, and a third (33%) that they always or frequently acted automatically and never or rarely intentionally (Fig. 1G and H). Like the false belief, which can range from vague (“something is off”) to specific (“a bomb is about to explode in the house”), the behavior can be directed towards a more or less specific goal. Sometimes patients describe ‘waking up’ with a sense of imminent danger and an urge to run, although they do not know what exactly is about to happen and what they are running away from. When the belief is more specific, that is, when patients ‘know’ what is going to happen, their acts usually reflect attempts to prevent the threat or its consequences, like hiding from an intruder, or running to save close ones from imminent danger. Although to outside observers, the behavior displayed during episodes may appear out of character and inappropriate, in these cases patients report that their actions are perfectly congruent with respect to the belief they hold during the episodes (Fig. 1H and I, Table 1 reports 16 – 22, 33 and 34). Purposeful behaviors can be highly skilled and complex. Examples reported by patients include writing grammatically correct text messages, speaking in a second language, dressing, correctly choosing between distance and reading glasses, remembering a previously established sequence of hand movements, or seeking out objects in the correct location and using them appropriately to perform a specific action (Table 1 report 34).

Almost all patients (94%) said they could interact with their environment (Fig. 1J); and 64% often or always did so (Fig. 1J). Not infrequently, patients involved their close ones or pets in their behavior. They reported dragging partners out of their bed to save them from natural disasters, verifying that their child was well, evacuating the house because a bomb was about to explode, or confronting their partner about their presumed infidelity. One patient recalled an episode during which he sat on his wife’s head to avoid a “flood of blue paint” from entering her mouth, almost suffocating her in the process. Another patient who frequently experienced choking on imaginary objects during his episodes started performing cardio-pulmonary massage on his daughter because he believed *she* had choked on something.

##### 6) Emotions

Most patients (75%) stated that their experiences during episodes were often or always unpleasant (Fig. 1K) and never or rarely pleasant, and roughly the same proportion (78%) characterized them as nightmares (Fig. 1L). 77% of patients reported never having had emotionally neutral experiences during episodes (Fig 1M) and 65% never having had pleasant experiences (Fig. 1L). Patients almost invariably described fear or apprehension as predominant emotions, and some patients even reported a distinctive feeling of imminent death (Table 1 report 35). Patients with sleep terrors sometimes pointed out that the intensity of the fear they experienced during their parasomnia episodes was incomparable to any waking life event (Table 1, reports 36 and 37).

##### 7) Other aspects

As can be seen in the reports presented in this paper (Table 1), patients often referred to their parasomnia episodes as dreams. When asked how these dreams compared to ‘ordinary dreams’, they frequently responded that the oneiric scenario associated with parasomnia episodes consisted in a single scene, without a narrative and changes of setting are typical of other dreams. Indeed, 69% of patients reported that experiences always or often consisted in a short single scene and were never or rarely story-like (Fig. 1O). For some patients, this scene ‘appeared out of nothing’, they could not tell what happened before, although some had the distinct impression that the scene was part of a longer dream that they forgot. Others reported that their episodes were triggered by surprising elements occurring during an otherwise ordinary dream (Table 1 report 14). Some patients referred to their parasomnia experiences as ‘little dreams’ to distinguish them from regular dreams. A minority of patients (22%) reported that experiences were always or often long and story-like and never short and scene-like (Fig 1P). Many patients found parasomnia experiences much more realistic and emotionally charged than other dreams. Some pointed out that during parasomnia episodes they were more active and felt the ‘urge to do something’, contrary to the more passive ordinary dreams. In addition, as opposed to ‘normal’ dreams, they sometimes perceived elements of their real surroundings, which gave them the impression of being awake. Indeed, although patients reported dreams, many, especially those with sleep terrors, also reported that they felt awake during episodes (referring to their episodes as ‘awake dreaming’). Interestingly, even when patients felt awake and experienced very intense fear during episodes, they could fall back asleep very quickly, sometimes in a matter of seconds (Table 1 report 38), suggesting a dissociation between emotional intensity and vigilance.

#### Parasomnia episodes with no experience (automatic behavior) or amnesia

38% of patients reported that they never or only rarely remembered their episodes (Fig. 1A). This finding inevitably raises the question of whether patients were amnestic of episodes, or whether such reports reflect a state of unconsciousness or minimal consciousness, characterized by behaviors without or little awareness. Although anecdotal reports of patients do not allow to draw definite conclusions, taken together, they suggest that *both* the level of consciousness and amnesia can be variable, not only between episodes in the same patient, but also within the same episode.

For instance, many patients described that they did not spontaneously recall their nocturnal episodes upon awakening in the morning, but could so later, when they noticed something unusual that reminded them of their nocturnal activity, or when witnesses described the episodes to them (Table 1 report 39). The fact that patients could remember their subjective experiences when provided with a clue suggests that the experience was correctly encoded in memory, although not spontaneously retrieved upon awakening. Among other factors that increase the likelihood of remembering episodes, patients mentioned being awakened by another person during the episode and negative emotions (‘I never seem to remember the gentle episodes, only the frightening ones’).

At other times, patients could describe what they were experiencing to their bedpartners during the episodes, although the next day they had no recollection of it. For instance, one patient who could not remember a single experience associated with his parasomnia episodes once suddenly jumped out of bed, woke up his wife and told her there were insects in his bed, pointing to the bedsheets and urging her to get out of bed. The next day, he did not remember the episode, even when his wife told him about it. Examples like this one, in which patients report their experiences to another person during the episode, and display behavior in line with their accounts, but do not recall them, strongly suggest that some episodes are associated with conscious experiences that are not encoded into memory.

Finally, there are episodes in which patients appear to display relatively automatic behaviors with no or little consciousness. Of course, most of these episodes cannot, by definition, be recounted in an interview. However, 81% of patients reported regaining consciousness while carrying out a particular activity at least once, with no specific intention or reason in mind ane 36% reported often or always having such episodes with automatic behavior (Fig 1G). These activities included gesticulating (Table 1 report 40), producing apparently senseless speech (Table 1 report 41), reflexively hitting another person who was approaching, or screaming without being aware of it (Table 1 report 42). One patient who presented short confusional arousals with a terrified facial expression every time his wife opened the squeaking bedroom door reported not experiencing any fear at all and regained consciousness only a few seconds later, when he saw his wife next to him. An anecdote provided by another patient (Table 1 report 42), suggests that awareness may be selective for some aspects of the experience. In this case, the patient did not hear his own screams, but was aware of his stepdaughter who was urging him to stop, and consequently, did not understand her request.

## Discussion

### Summary of findings

The results of this study suggest that consciousness during Non-REM parasomnia episodes is variable: some episodes appear to have occurred without apparent consciousness and largely automatic behaviors, while others were associated with vivid conscious experiences characterized by complex thinking, realistic multisensory hallucinations as well as skilled, highly purposeful behaviors and interactions with the environment. This variability was present both between as well as well as within individuals (across episodes). Although the degree of amnesia for parasomnia episodes was overall high, it also featured some variability: certain episodes were vividly remembered, others could only be remembered when patients were provided with appropriate cues, and still others did not seem to be encoded in memory at all. Despite this variability in consciousness and amnesia, when episodes were remembered, they displayed several invariable features *(Table 2)*: they were characterized by delusional thoughts, mostly about impending danger or death, as well as negative emotions (fear, apprehension), with behaviors reflecting attempts to prevent the imaginary threat or its consequences. Delusional thoughts were almost never criticized, although doubts and ambivalence sometimes occurred towards the end of the episodes.

Similar to the *level* of consciousness, which was present to various degrees, there was also a gradation in the representation of the *contents* of consciousness (Table 2 and Fig. S1). Thoughts could range from vague ideas of threat to specific and elaborated delusions. Sensory experiences (hallucinations) encompassed unstructured shadows and silhouettes on one extreme, and highly realistic representations of objects, living beings or places on the other. The degree to which sensory experiences reflected the current environment was also graded: experiences could be entirely unrelated (hallucinations), partially related (misidentifications of people, perceptual illusions of objects) or presumably adequately reflect the current environment (perception), although this latter aspect is more difficult to ascertain a posteriori. Thus, the sensory disconnection that characterizes sleep appears to be variably present during NREM parasomnia episodes.

### Comparison with previous studies

The core features of NREM parasomnia experiences described herein are in line with previous work. Studies including adults with NREM parasomnias have described high recall rates of episodes over a lifetime (67-91% ^13,15,18,21^), mostly distressing and threatening mental contents (64-70% ^13,25^), predominantly negative emotions (84%^18^), a greater amount of single scene over story-like dream experiences (94%^9,18^) and the frequent setting of the dreams within the current environment (42 %^13^). Descriptions of delusions and hallucinations figure in several case reports ^5,12,14,16,19,20,26^ and larger studies ^9,13,17,18,27^. Sleepwalking was long conceptualized as preprogrammed ambulatory automatism that once it has started, completes its course ^6,28^, and unresponsiveness to the environment was initially described as the rule. Later studies emphasized goal-directed behaviors in line with the dream contents ^5,13,18,29^, giving the impression that NREM parasomnias, as a rule, reflect dream enactment. The experiences described herein provide a more nuanced view, suggesting that consciousness is variable and graded across individuals and episodes, similar to the variability in the level of consciousness that sleeping individuals report upon awakening from NREM sleep^10,30–32^, ranging from the absence of experience to full-fledged dreams. They also allow to qualitatively refine and complement previous descriptions of parasomnia experiences, showing that both encoding and retrieval deficits for memories of parasomnia episodes can occur, that hallucinations can be multisensory in nature and appear to be variably formed, that the environment is experienced on a perception-illusion-hallucination continuum, that patients frequently feel awake during episodes and that they have profoundly limited insight and reality monitoring despite being able to perform risk calculations and displaying perfectly intentional behaviors.

### Parasomnia experiences vs. ‘ordinary’ dreams

Many patients in this study spontaneously described their parasomnia experiences as dreams or nightmares. Similar to typical dreams, parasomnia experiences are characterized by delusional aspects (false beliefs), a lack of lucidity and control over the experience, multisensory (but mainly visual) hallucinations, heightened emotionality, disorientation in time and variable, but often pronounced amnesia^3,33,34^. Experiences reported by patients were short and mostly consisted in a single scene, which could indicate that conscious experiences were indeed shorter, but also attentional reorientation within a dream, or the creation of a new dream scene. In this respect, Non-REM parasomnia experiences appear more similar to Non-REM dreams, which are generally shorter and more realistic^35–37^ compared to REM dreams, although they may, at times, be longer and remarkably vivid^33^. These similarities between Non-REM parasomnia experiences and dreams are supported by a recent laboratory study, in which Non-REM parasomnia episodes were induced with loud sounds during slow wave sleep, and patients were interviewed immediately afterwards about their experiences^38^. Compared to episodes for which patients reported no experience (unconsciousness, 19%), those with report of conscious experiences (81%) were preceded by a higher degree of regional cortical activation prior to movement onset, similar to brain activity patterns that distinguish dreams from no experience in both Non-REM and REM sleep^31,39^. In addition, episodes with unconsciousness tended to be short and feature relatively stereotyped arousal-related behaviors, while those with dream-like experiences had a more variable length. They featured a similar stereotyped beginning, but then unfolded in a more idiosyncratic way. It is thus conceivable that a sudden activation of arousal systems, as the one induced by loud sounds in the study, is secondarily contextualized as a dream (or within a dream) only when the brain is in a state to do so, i.e., the cortex is regionally activated. A stereotyped activation of arousal systems could also explain the relatively stereotyped dream contents reported in the present study, consistently revolving around impending threat. Remarkably, the specific contents of delusional thoughts and hallucinations were not only reported by different patients in this study, but also in the literature ^11,13,18^. These similarities between parasomnia episodes deserve further consideration and systematic comparison with other dreams. Another major difference to ordinary dreaming is that during parasomnia episodes, patients were not disconnected from their surroundings, they showed overt behaviors and could also perceive environmental elements.

### Beyond ordinary dreams

The experiences reported in this study illustrate how during sleep-related states, normally integrated aspects of consciousness, including memory, reflective consciousness, control over behavior and the perception of the environment can selectively disintegrate: behavior can become dissociated from consciousness, implausible ideas from critical discernment, emotional intensity from vigilance and sensory experiences from environmental stimuli. While some of these dissociations are also seen in dreams^40^, the study of parasomnia episodes is particularly instructive with regards to how the brain creates a representation of reality. The observations reported herein suggest that individuals with Non-REM parasomnias ‘wake up’ with a distinct expectation of danger that not only determines what they do, but also what they perceive: well-meaning bedpartners are taken for enemies, a cloth rack is seen as a dangerous power line with bats on it, and straps of a bag mistaken for snakes. In other words, during parasomnia episodes, patients see what they believe and believe what they see. Some reports of patients suggest that degree of sensory ‘evidence’ that is present at a given time could constrain the possibilities of what is experienced. For instance, in some cases, ambient illumination or the presence of other sensory stimuli (i.e. bedpartners talking) favored the perception of the real environment or illusions and misidentifications over full-fledged hallucinations. A similar effect has been experimentally demonstrated for mental imagery, which is attenuated in conditions of high luminance ^41^. Thus, sleepwalking could be an exemplary case, in which internally and external generated ‘stimuli’ compete against each other to determine what is ultimately ‘experienced’.

### Limitations

The data presented herein, is, per study design (interviews), biased towards the description of episodes with conscious experiences. Episodes without recall or with mainly automatic behaviors can only be indirectly inferred, and it is possible that they are more frequent than reported herein. The exclusion of children^21^ and of patients presenting exclusively with sexsomnia or sleep-related eating, who are often not conscious during their episodes, may also have biased results in this direction. However, the absence of experience (unconsciousness) was reported only after 19% of Non-REM parasomnia episodes by adult patients who were interviewed immediately after the episode in the laboratory, suggesting that they are indeed the minority^38^. Some features of conscious experiences could make them more prone to be remembered, including, as reported by several patients, the presence of fear. Indeed, expressions of fear were only seen in 23% of episodes in a laboratory study^38^. Thus, it is possible that the features that were identified as relatively invariable in this study apply only to a specific subset of episodes that are preferentially remembered in the long term. Finally, the limited sample size, albeit comparable to other studies of this type ^13,15,18^, did not allow for subgroup analyses taking age, parasomnia subtype, and other factors into account. Information in this study was collected during in-depth exchanges with patients, lasting in some cases up to two hours, which was crucial to understand what patients experienced, obtain many examples, and make sure participants understood the statements they rated. However, this procedure also limited the number of patients that could be included. In the future, it would be highly valuable to perform such a study in a larger population.

## Data availability statement

A large part of the data (reports, etc.) is directly presented in the article. Additional information or data is available upon request.

## Supplementary Material

**Figure S1:**
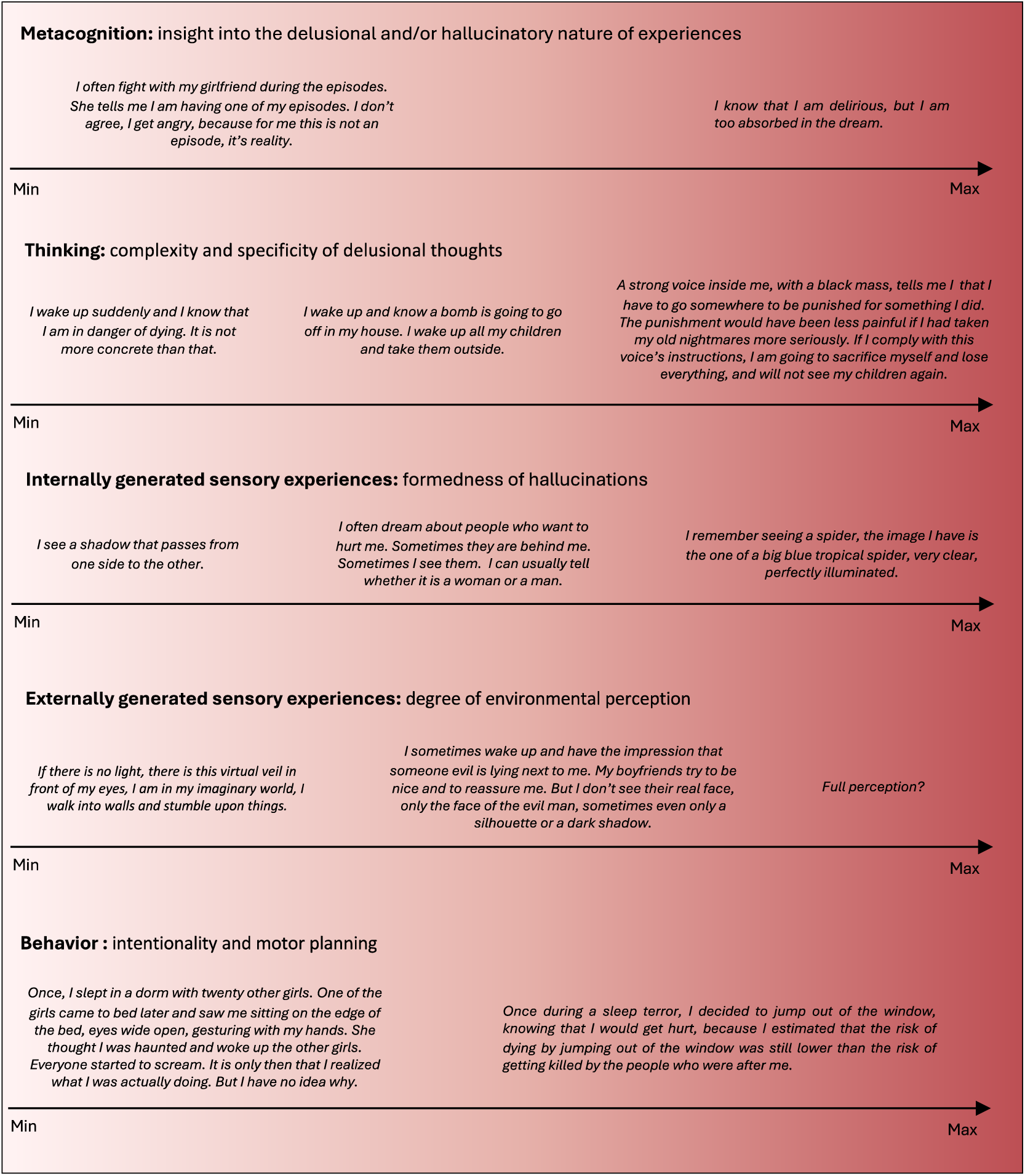
The spectrum of cognitive dimensions in parasomnia experiences illustrated with reports of patients.

